# RGF1 controls PLT2 protein stability through ROS-dependent regulation of a cysteine residue in root meristem development

**DOI:** 10.1101/2024.04.08.588570

**Authors:** Yu-Chun Hsiao, Shiau-Yu Shiue, Ming-Ren Yen, Joon-Keat Lai, Masashi Yamada

## Abstract

The protein concentration gradients of the master regulators of the root meristem, named the PLETHORA proteins, modulate the root meristem size. Root meristem growth factor 1 (RGF1) peptide extends the PLETHORA2 (PLT2) protein gradients by altering reactive oxygen species (ROS) distributions. However, the underlying mechanism through which the ROS alterations regulate PLT2 remains unknown. Here, we demonstrate that the 212^th^ cysteine of the PLT2 protein plays a pivotal role in modulating PLT2 stability through the ROS altered by RGF1. The substitution of the 212^th^ cysteine of PLT2 with serine (PLT2^C212S^) enhanced the PLT2 protein stability upon RGF1 and resulted in robust resistance to ROS relative to the native PLT2. Accordingly, PLT2^C212S^ modulated expressions of certain specific root development-related genes to a greater extent than native PLT2. In summary, these findings show that the PLT2 concentration gradient formation through ROS, modulated by RGF1, is dependent on a mechanism involving the 212^th^ cysteine of PLT2.

## Text

As sessile organisms, plants grow and develop organs in response to various environmental changes. Therefore, plants require real-time, sensitive cell-to-cell communications to control their development and adapt. Plant meristems enable plants to develop and produce new organs throughout their lifespan to optimize their growth. Plant roots show dynamic growth, and this process is tightly coordinated by each root zonation, i.e., the meristematic, elongation, and differentiation zones^1^. Stem cell daughters undergo active cell divisions in the meristematic zone to produce numerous small cells; after that, cells rapidly expand to increase their size and finally differentiate to acquire distinct structures^1^. The size of each developmental zone is determined by phytohormones, such as auxin and cytokinin, as well as master regulators, PLETHORAs (PLTs), which are critical for maintaining root stem cell activity and meristem growth^2^. PLT members contain two conserved *APETALA2*/*ETHYLENE-RESPONSE FACTOR* (*AP2/ERF*) DNA binding domains^3,4^. The transcripts and promoter activities of the *PLT1*, *PLT2*, *PLT3*, and *BABY BOOM* (also known as *PLT4*) genes are detected around the quiescent center (QC)^3,4^. However, the proteins of PLTs display broader gradient distributions and partly overlap in the meristematic zone^4^, suggesting coordinated post-translational regulation defines their gradients. In addition, they function redundantly in stem cell maintenance.

PLT1 and PLT2 play particularly predominant roles in the root meristem^3,4^. PLTs dose-dependently control developmental events in the root developmental zones, suggesting the sophisticated PLT gradient defines root zonation^4^.

The sulfated ROOT MERISTEM GROWTH FACTOR (hereafter referred to as RGF, also known as GOLVEN or CLE-like) peptides also play pivotal roles in maintaining the root meristem and determining the root meristem size^6–8^. The *rgf* single mutant does not show significant defects in the root, whereas the *rgf1/2/3* triple mutant reduces the root meristem size^6^. External supply with sulfated synthetic RGF1 peptide rescues the *rgf1/2/3* root meristem defects, showing that the RGF1 dose-dependently functions in root meristem development^6^. Our recent work showed that treatment with RGF1 expanded PLT2 protein localization by increasing superoxide (O_2_^-^) in the meristematic zone and decreasing hydrogen peroxide (H_2_O_2_) levels around the boundary between the meristematic and elongation zones^9^. The broadly localized PLT2 formed a larger root meristem^9^. Co-treatment of RGF1 with an NADPH oxidase inhibitor, which reduces O_2_^-^, or a high dose H_2_O_2_ (strong root growth inhibition), reduced the extended PLT2 localization^9^. In contrast, co-treatment with RGF1 and an H_2_O_2_ scavenger, which reduces H_2_O_2,_ extended the PLT2 protein localization relative to the RGF1-only treatment^9^. These findings suggested that the increased O_2_^-^ and reduced H_2_O_2_ levels modulated by RGF1 extended the localization of the PLT2 protein. RGF1 treatment increased the PLT2 protein stability but not the *PLT2* promoter activity^9^. These findings suggested that the ROS distributions modulated by RGF1 post-translationally alter the PLT2 protein stability. However, the underlying molecular mechanism of how the ROS altered upon RGF1 affects the PLT2 protein stability in the root meristem remained obscure in these works.

Here, we show that the 212^th^ cysteine residue of the PLT2 protein predominantly affects ROS-dependent PLT2 stability by RGF1. Replacement of the 212^th^ cysteine with serine (PLT2^C212S^) of PLT2 resulted in a broader gradient and higher amount of PLT2 with a larger meristematic zone upon RGF1 treatment and resistance to ROS, in comparison to replacement with another cysteine (PLT2^C112S^) and native form PLT2. Furthermore, the PLT2^C212S^ treated with RGF1 modulated the expression of a similar array of genes to native PLT2, but to a greater extent. Upon RGF1 treatment, PLT2^C212S^ upregulated genes associated with cell proliferation expressed in the meristematic zone while simultaneously downregulating those genes linked to cell maturation localized in the elongation zone. Our results, thus, clearly show the post-translational regulation mechanisms of PLT2 protein stability via ROS upon RGF1 treatment, and the PLT2 downstream gene expressions under RGF1 signaling.

## Results

### PLT2 is a critical regulator in the RGF1 signaling pathway

The *plt1/2/3* mutants have a redundant phenotype of root meristem maintenance^4^. To understand which *PLT* is involved in RGF1 signaling, we examined the response of individual *plt* mutants to RGF1 peptide. We monitored the wild-type, *plt1*, *plt2*, and *plt3* root phenotypes under the confocal microscope after 5 nM RGF1 treatment for 24 h. The meristematic zone size was dramatically increased in the wild-type, *plt1*, and *plt3* after adding RGF1; however, the *plt2* mutant was almost insensitive to the peptide treatment with a meristematic zone size comparable to that of the Mock treatment (Fig. 1a, b), indicating that PLT2 is the dominant regulator defining the meristematic zone size in the RGF1 signaling pathway.

**Figure 1.**
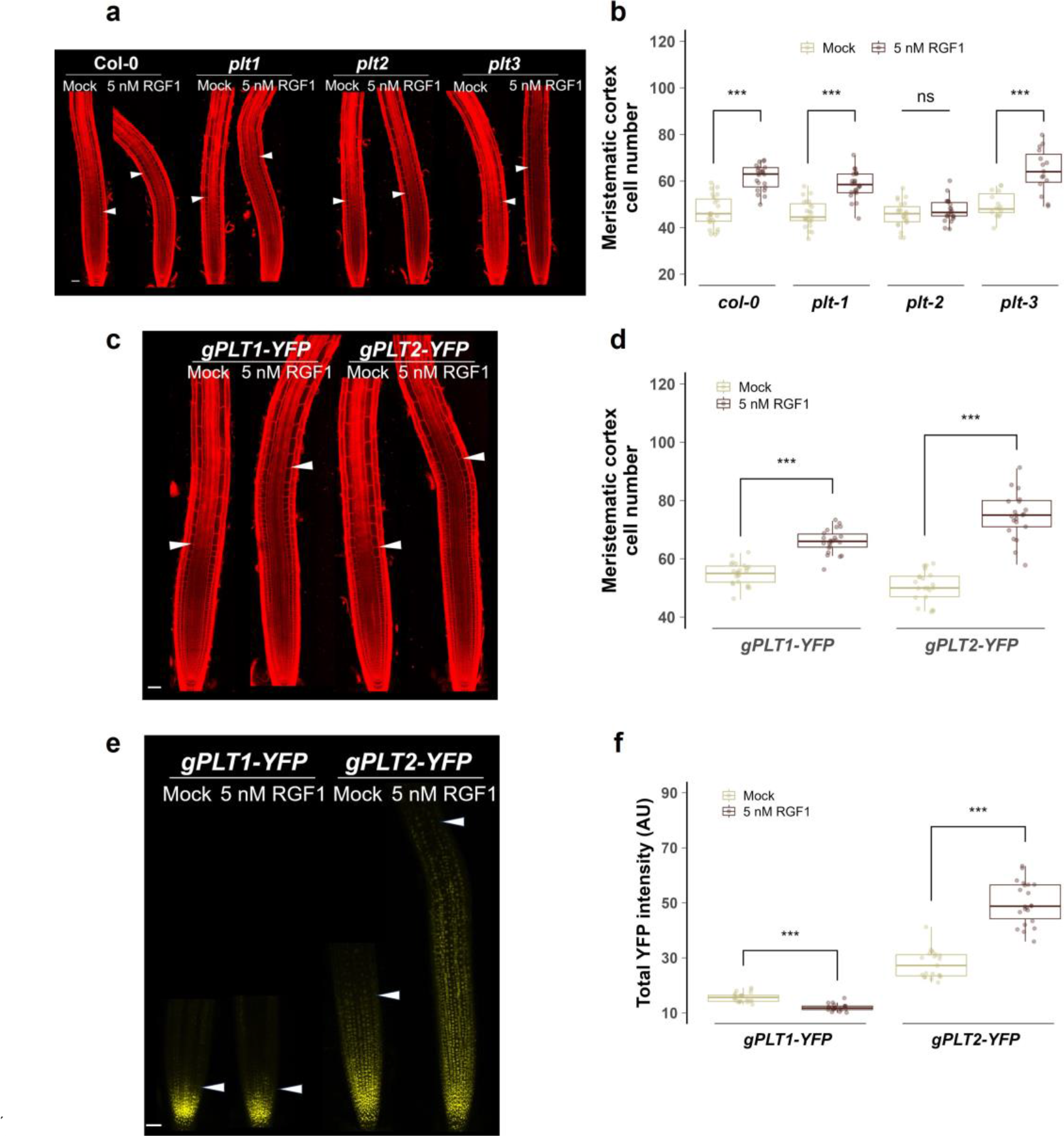
PLT2 protein is a critical regulator in the RGF1 signaling pathway. **a**, Confocal images of *Col-0*, *plt1*, *plt2*, and *plt3* roots after mock or 5 nM RGF1 treatment for 24 h followed by PI staining. The white arrowheads show the junction between the meristematic and elongation zones. **b**, Number of cells in the meristematic zone in *Col-0*, *plt1*, *plt2*, and *plt3* roots after mock or 5 nM RGF1 treatment for 24 h. **c**, **d**, **e**, **f**, Confocal images (c and e) and the quantification analysis (d and f) of *gPLT1-YFP/Col-0* and *gPLT2-YFP/Col-0* roots after mock or 5 nM RGF1 treatment for 24 h. The white arrowheads show the junction between the meristematic and elongation zones, and in (e) show the extent of YFP expression. In **b**, **d**, and **f**, the boxplots show the individual data points (circles), median (horizontal line within boxes), interquartile range (IQR) (hinges), and the 1.5 * IQR extension (whiskers). In the Student’s t-test in b, d, and f, the symbols *** and ns respectively indicate a P value less than 0.001 and greater than 0.05. All scale bars, 50 μm.

The protein sequence of PLT2 is highly similar to that of PLT1^4^. We next investigated how RGF1 regulates the PLT1 and PLT 2 proteins by observing the protein localizations of gPLT1-yellow fluorescent protein (YFP) and gPLT2-YFP^4^ in the wild-type background upon supplying RGF1. The root meristem size was increased in both genetic backgrounds by RGF1, showing that RGF1 treatment works in both lines (Fig. 1c, d).

The gPLT2-YFP signal showed extended localization after adding RGF1. In contrast, the gPLT1-YFP signal was restricted around the root apical meristem (RAM) irrespective of the RGF1 supplement (Fig. 1e). In addition, we found the gPLT1-YFP signal was slightly reduced after RGF1 treatment (Fig. 1e, f), suggesting the distinct roles of PLT1 and PLT2 in regulating root meristem development under the RGF1 signaling pathway. This data again confirmed that PLT2 is the core component defining the meristem size in the RGF1 pathway.

### The Cys residue is essential for the modulation of PLT2 protein stability by RGF1

ROS altered by RGF1 modulate the PLT2 protein stability^9^. ROS are known to modify specific cysteine (Cys) residues to regulate the protein stability and activity in various biological and physiological processes^10,11^ Two Cys residues appear among the PLT2 sequences, the 112^th^ and 212^th^ amino acids, and the latter residue, found in the *AP2/ERF* domain, is conserved in PLT1 and PLT2 (Extended Data Fig. 1). Therefore, we hypothesized that the Cys residues of PLT2 may be essential for the protein stability modulated by ROS. To test the hypothesis, we replaced one or both of the Cys residues of PLT2 with serine (Ser) by site-directed mutagenesis. We replaced Cys with Ser because the difference between Cys and Ser is only the substitution of one sulfur atom with oxygen. The constructs of the single Cys substitution genomic *PLT2^C112S^* (*gPLT2^C112S^*), *gPLT2^C212S^*, and the double Cys substitution (*gPLT2^C112SC212S^*), as well as the native *gPLT2* (hereafter designated as the wild-type *PLT2*) fused with green fluorescent protein (*GFP*) driven by the *PLT2* promoter, were introduced in the *plt2* mutant background for further analysis. Indeed, all the introduced constructs of *gPLT2* with or without Cys replacement rescued the insensitivity to RGF1 of the *plt2* mutant (Fig. 2a, b). Notably, the meristem sizes and cell numbers of the 212^th^ Cys replacement line were substantially increased by RGF1 compared to those of the wild-type and the 112th Cys substitution lines (Fig. 2a, b). However, the phenotypes of the *gPLT2^C112S^-GFP* line were similarly increased by RGF1 compared to those of the wild-type PLT2 (Fig. 2a, b). The 112^th^ and 212^th^ double Cys substitution line showed an equivalent potent response upon RGF1 treatment in comparison with the 212^th^ single Cys substitution line, suggesting that the 112^th^ Cys does not have an additive effect (Fig. 2b). These results indicated that the 212^th^ Cys contributes to the expansion of the meristematic zone size with RGF1 treatment.

**Figure 2.**
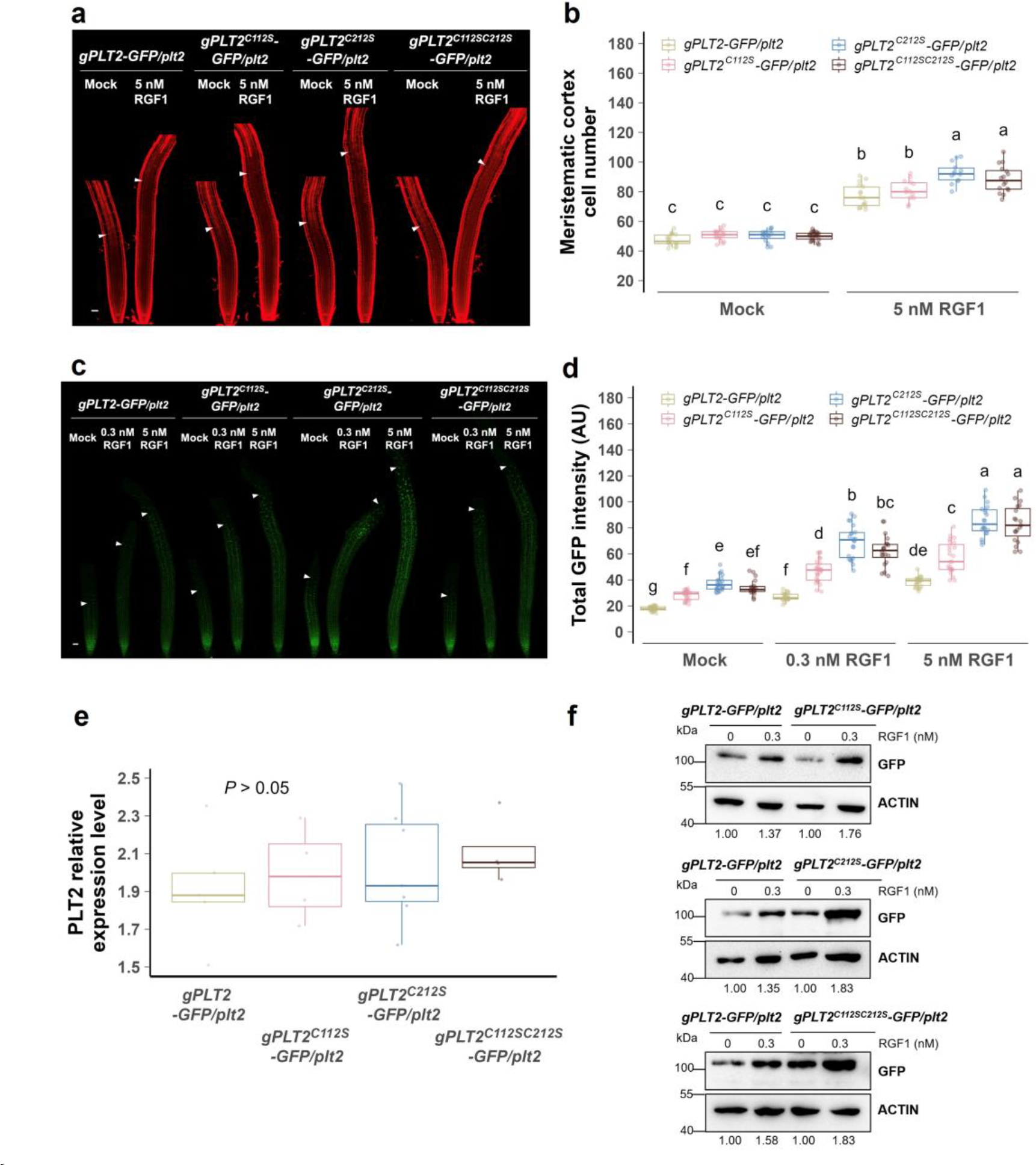
The cysteine residue of the PLT2 protein is essential for the RGF1-modulated PLT2 stability. **a**, **b**, Confocal images (a) and the meristematic cell number (b) of *gPLT2-GFP*/*plt2*, *gPLT2^C112S^-GFP*/*plt2*, *gPLT2^C2^*^12^*-GFP*/*plt2*, and *gPLT2^C112SC212S^-GFP*/*plt2* roots after mock or 5 nM RGF1 treatment for 24 h. Propidium iodide (PI) staining is in red. The white arrowheads indicate the junction between the meristematic and elongation zones. **c**, **d**, Confocal images (c) and the quantification of total GFP intensity (d) in the meristematic zone of *gPLT2-GFP*/*plt2*, *gPLT2^C112S^-GFP*/*plt2*, *gPLT2^C212S^-GFP*/*plt2*, and *gPLT2^C112SC212S^-GFP*/*plt2* roots treated with either mock, 0.3 nM, or 5 nM RGF1 for 24 h. The white arrowheads indicate the extent of GFP expression. **e**, qRT-PCR quantification of the *PLT2* expression in three developmental zones of *gPLT2-GFP*/*plt2*, *gPLT2^C112S^-GFP*/*plt2*, *gPLT2^C212S^-GFP*/*plt2*, and *gPLT2^C112SC212S^-GFP*/*plt2* after either mock or 0.3 nM RGF1 treatment for 24 h. **f**, Western blot analysis of PLT2 protein in *gPLT2-GFP*/*plt2*, *gPLT2^C112S^-GFP*/*plt2*, *gPLT2^C212S^-GFP*/*plt2*, and *gPLT2^C112SC212S^-GFP*/*plt2* roots after either mock or 0.3 nM RGF1 treatment for 24 h. ACTIN was the loading control. GFP normalized with ACTIN ratio is shown at the bottom of the graphs. In **b**, **d**, and **e**, the boxplots show the individual data points (circles), median (horizontal line within boxes), interquartile range (IQR) (hinges), and the 1.5 * IQR extension (whiskers). Two-way ANOVA with Tukey’s post hoc test in **b** and **d** and one-way ANOVA in **e**. Boxes not sharing a lower-case letter indicate a significant difference (*P* < 0.05). Scale bar, 50 μm (a, c).

In this study, we determined middle (0.3 nM) and high (5 nM) concentrations of RGF1 based on the induced O_2_^-^ levels. We detected the O_2_^-^ level in each PLT2 transgenic line by nitro blue tetrazolium (NBT) staining using treatments with these two different RGF1 concentrations. Correspondingly, in all PLT2 transgenic plants, RGF1 applications dose-dependently induced O_2_^-^ accumulation in the meristematic zone; however, there were no significant differences across genetic backgrounds (Extended Data Fig. 2a, b), showing that RGF1 modulates ROS levels equivalently in all genetic backgrounds.

Next, we examined how the ROS altered by these two concentrations of RGF1 affect the PLT2 stability via the Cys. We found that the localizations of the PLT2^C112S^-GFP were modestly increased by each application of RGF1 relative to the wild-type PLT2-GFP (Fig. 2c, d). Notably, the PLT2^C212S^-GFP and PLT2^C112SC212S^-GFP proteins displayed broader localization and higher intensities than the wild-type PLT2-GFP and PLT2^C112S^-GFP upon each RGF1 treatment (Fig. 2c, d). These results indicated that the 212^th^ Cys residue is pivotal for the PLT2 stability in the RGF1 signaling pathway. A similar trend was observed in the second independent transgenic line of each PLT2 construct (Extended Data Fig. 3a-d).

Furthermore, we conducted an immunoblot assay to examine the PLT2 protein amount after 0.3 nM RGF1 treatment by collecting the root tip tissues, including three developmental zones in each PLT2 transgenic plant, because O_2_^-^ levels are still not saturated at 0.3 nM RGF1 (Extended Data Fig. 2a, b). The ratio of the GFP protein levels relative to the ACTIN (ACT) protein by the RGF1 treatment in the gPLT2^C112S^-GFP line was increased to a moderately greater extent than that of the gPLT2-GFP line (Fig. 2f, under the panel). However, the fold change of GFP relative to ACT was elevated in the gPLT2^C212S^-GFP and gPLT2^C112SC212S^-GFP lines to a higher degree than the other two lines upon RGF1 treatment. These results confirmed that the 212^th^ Cys residue plays a vital role in the regulation of PLT2 protein stability by RGF1.

ROS levels were similarly increased according to the concentration of the RGF1 treatment in the wild-type PLT2 and all Cys substitution lines (Extended Data Fig. 2a, b). The four constructs had similar gene structures except for the sequences of one or two cysteines. These data thus suggested that RGF1 modulated PLT2 protein stability via the 212^th^ Cys by ROS-dependent post-translational regulation. To clarify this, we examined *PLT2* expression in each PLT2 genetic background using quantitative reverse transcription polymerase chain reaction (qRT-PCR). The *PLT2* transcript increase ratio was comparable in all transgenic plants after the peptide treatment (Fig. 2e), suggesting that the broader localization of PLT2 protein in the 212^th^ Cys substitution lines by RGF1 was caused by ROS-dependent post-translational regulation but not transcriptional regulation.

Since it has been reported that the PLT2 protein moves from cell to cell^13^, RGF1 may induce PLT2 protein movement. To ascertain whether the extended localization of PLT2 after RGF1 treatment was due to protein transfer, the construct of the wild-type *PLT2* fused to 3x*GFP* driven by the *PLT2* promoter was introduced in the *plt2* mutant background because the high molecular weight of the 3xGFP tag reduces the protein movement^14^. After RGF1 treatment, the PLT2-3xGFP protein was broadly localized at an equivalent level to the PLT2-1xGFP protein distribution (Extended Data Fig. 4a, b), indicating that the broader localization of the PLT2-3xGFP protein was more likely to be enhanced by the PLT2 protein stability rather than the protein transportation. Furthermore, immunoblot assay (Fig. 2f) showed that the protein amounts were highly elevated by RGF1 treatment in all PLT2 transgenic lines, including the Cys substitution lines, suggesting that RGF1 regulates the PLT2 protein stability via ROS rather than protein movement.

### ROS levels control the PLT2 protein stability mainly via the 212^th^ Cys

We quantified the GFP intensities of PLT2 in the epidermal cells in each represented developmental zone, which were around the apical root meristem, the proximal side (close to the elongation zone) of the meristematic zone, and the elongation zone (Fig. 3a, red, yellow, blue arrowheads), in the wild-type *PLT2* and the *gPLT2^C212S^-GFP* lines. The GFP signal in the wild-type PLT2 line decreased rapidly and became dim on the proximal side-meristematic and elongation zones, where H_2_O_2_ is accumulated^9^ (Fig. 3a, b). The PLT2^C212S^ protein was more stable and broadly detected in these two developmental zones, thus showing a relatively strong signal in the cell expansion area (Fig. 3a, b) with a higher chance of contact with H_2_O_2_^9^. Therefore, we hypothesized that H_2_O_2_ reduced the wild-type PLT2 protein level around the elongation zone; however, PLT2^C212S^ was not susceptible to H_2_O_2_ and was still stable even at the cell elongation area. To test our hypothesis, we exogenously applied a high dose (1 mM, strong inhibition of root growth) of H_2_O_2_ for 24 h and observed how the roots of each genetic background behaved under the conditions. The meristem size of the wild-type PLT2 was significantly reduced after a high concentration of H_2_O_2_ treatment (Fig. 3c, d). The root meristem size of *gPLT2^C112S^-GFP* showed a mild reduction after H_2_O_2_ treatment(Fig. 3c, d). The wild-type PLT2-GFP and the PLT2^C112S^-GFP signals were markedly decreased after H_2_O_2_ application (Fig. 3e, f). Surprisingly, the *gPLT2^C212S^-GFP* and *gPLT2^C112SC212S^-GFP* lines demonstrated a robust resistance to H_2_O_2_, and the roots were intact and showed similar GFP intensities to the Mock treatment (Fig. 3c-f), suggesting that H_2_O_2_ mainly targets the 212^th^ Cys to modulate PLT2 protein stability. Furthermore, we confirmed the confocal observation by immunoblot analysis. The wild-type PLT2 and PLT2^C112S^ proteins were labile when exposed to a high dose of H_2_O_2_, thus degrading rapidly (Fig. 3g). However, the PLT2^C212S^ and PLT2^C112SC212S^ proteins were stable, albeit under high H_2_O_2_ conditions (Fig. 3g). These results revealed that the wild-type PLT2 protein was susceptible to H_2_O_2_, PLT2^C112S^ showed a mild resistant response, and PLT2^C212S^ was almost insusceptible to H_2_O_2_. We did not observe an additive effect in the PLT2 line with a double Cys replacement; therefore, we concluded that H_2_O_2_ was triggered to control the PLT2 protein stability predominantly on the 212^th^ Cys.

**Figure 3.**
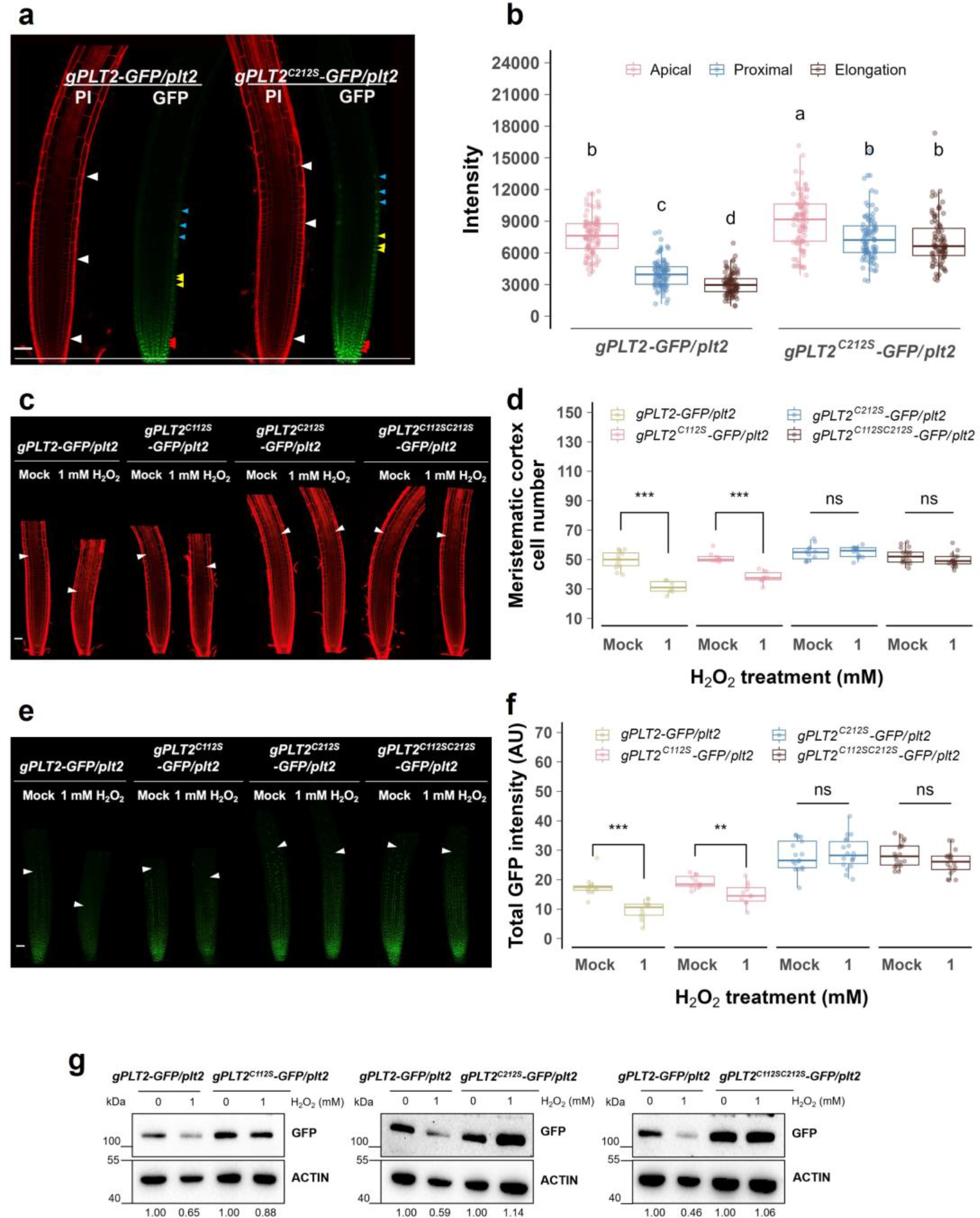
Redox status controls the PLT2 protein stability via the 212^th^ cysteine. **a**, The PLT2 protein localization in the epidermal cells in the apical root meristem (below the bottom white arrowheads close to the root tip), the proximal side (close to the elongation zone, below the middle white arrowheads) of the meristematic zone, and the elongation zone (between the top and the middle white arrowheads) in *gPLT2-GFP*/*plt2* and *gPLT2^C212S^-GFP*/*plt2* lines. **b**, The quantification of GFP fluorescence per cell in the apical root meristem (a, red arrowheads), the proximal side of the meristematic zone (a, yellow arrowheads), and the elongation zone (a, blue arrowheads) in *gPLT2-GFP*/*plt2* and *gPLT2^C212S^-GFP*/*plt2* lines. **c**, **d**, **e**, **f**, Confocal images (c, e) and the quantification analysis (d, f) of *gPLT2-GFP*/*plt2*, *gPLT2^C112S^-GFP*/*plt2*, *gPLT2^C212S^-GFP*/*plt2*, and *gPLT2^C112SC212S^-GFP*/*plt2* roots treated with either mock or 1 mM H_2_O_2_ for 24 h. The white arrowheads in (c and e) show the junction of the meristematic zone and the extent of GFP expression, respectively. **g**, Western blot analysis of PLT2 protein in *gPLT2-GFP*/*plt2*, *gPLT2^C112S^-GFP*/*plt2*, *gPLT2^C212S^-GFP*/*plt2*, and *gPLT2^C112SC212S^-GFP*/*plt2* roots after either mock or 1 mM H_2_O_2_ treatment for 24 h. ACTIN was the loading control. GFP normalized with ACTIN ratio is shown at the bottom of the graphs. In **b**, **d**, and **f**, the boxplots show the individual data points (circles), median (horizontal line within boxes), interquartile range (IQR) (hinges), and the 1.5 * IQR extension (whiskers). Two-way ANOVA with Tukey’s post hoc test in **b**. Boxes not sharing a lower-case letter indicate a significant difference (*P* < 0.05). Student’s t-test in **d** and **f**. The symbols *\*\*\**, *\*\**, and *ns* respectively indicate P value of less than 0.001, 0.01, and greater than 0.05. Scale bar, 50 μm (a, c, e).

Furthermore, to test the effect of O_2_^-^ on the PLT2 protein stability, we treated RGF1 with a low concentration (700 nM, minor root growth defects) of diphenyleneiodonium (DPI) to reduce O_2_^-^ levels (Extended Data Fig. 5a, b). Co-treatment with RGF1 and DPI reduced PLT2 stabilities in the wild-type PLT2 and *gPLT2^C112S^-GFP* lines markedly compared with RGF1 single treatment until they were a similar level to those of Mock treatment (Extended Data Fig. 5a, b). In contrast, the gPLT2^C212S^ and gPLT2^C112SC212S^ proteins were slightly reduced by the co-treatment but not diminished to the degree of the Mock treatment (Extended Data Fig. 5a, b), suggesting that lower O_2_^-^ levels affect the PLT2 stability via the 212^th^ Cys. The data from H_2_O_2_ and DPI treatments suggested that higher O_2_^-^ and H_2_O_2_ levels, respectively, stabilize and reduce the PLT2 protein through the 212^th^ Cys.

### PLT2^C212S^ modulates specific root development-related downstream genes after RGF1 treatment to a higher extent than wild-type PLT2

High levels of PLT control stem cell maintenance, moderate levels promote cell proliferation, and lower levels are required for cell differentiation, suggesting the sophisticated PLT gradient defines root development and zonation^4,13^ . Microscopic analysis showed that RGF1 treatment decelerated the PLT2 degradation, enlarged the meristematic zone size, and delayed the initiation of the cell elongation and differentiation in the *gPLT2^C212S^-GFP* line to a greater extent than the wild-type PLT2 line (Fig. 2a-d). Therefore, we hypothesized that a higher amount of PLT2^C212S^ affected more downstream genes required for the specific root developmental zones. To assess the impacts of PLT2 with or without the 212^th^ Cys in the *plt2* mutant at the transcriptome level, we performed RNA sequencing (RNA-seq) analysis in root tips containing the three developmental zones 24 h after mock, 0.3, and 5 nM RGF1 treatment. Differentially expressed gene (DEG) analysis was conducted for the *plt2* mutant, *gPLT2-GFP*/*plt2*, and *gPLT2^C212S^-GFP*/*plt2* (Fig. 4a). Interestingly, the *gPLT2^C212S^-GFP/plt2* had over 8,000 DEGs (encompassing both up and downregulated genes, 3,885 and 4,307, respectively) after 0.3 nM RGF1 treatment, which was much higher than the over 6,400 DEGs (2,995 up and 3,434 downregulated) in *gPLT2-GFP/plt2* even under 5 nM of the peptide supplement (Fig. 4a). In the comparison between *gPLT2-GFP/plt2* and *gPLT2^C212S^-GFP/plt2,* 3632 positively and 4032 negatively regulated DEG genes were overlapped (Fig. 4b, inside the intersection in the Venn diagram). These common up and downregulated DEGs were modulated to a greater extent in *gPLT2^C212S^-GFP/plt2* than in *gPLT2-GFP/plt2* by the RGF1 treatments (Fig. 4c). These results suggest that the additional accumulation of PLT2^C212S^ protein by RGF1 modulates downstream genes more strongly.

**Figure 4.**
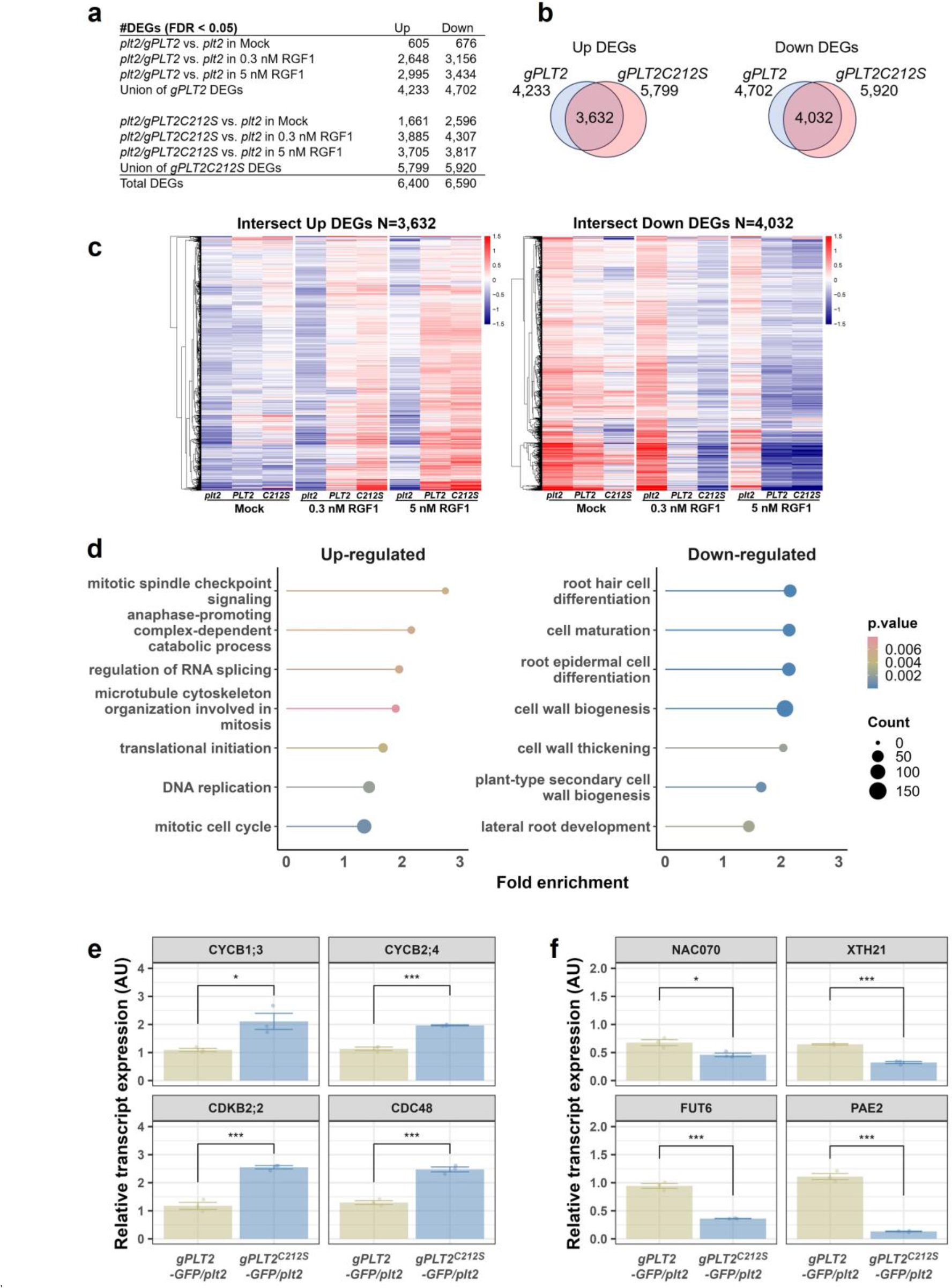
PLT2^C212S^ induces similar genes to PLT2 after RGF1 treatment but to a greater extent. **a**, The number of DEGs in the *gPLT2-GFP* and *gPLT2^C212S^-GFP* lines upon RGF1. **b**, Venn diagram showing the overlap of the up and downregulated DEGs in the *gPLT2-GFP* and *gPLT2^C212S^-GFP* lines. **c**, A heat map of the DEGs shared in the *gPLT2-GFP* and *gPLT2^C212S^-GFP* lines. **d**, The GO terms significantly enriched within the shared positively or negatively regulated DEGs after RGF1 treatment (Supplementary Data Set 2). In **e and f**, the bar plots show the individual data points (circles), the mean (bars), and the standard error (error bars). Student’s t-test in **e** and **f**. The symbols *\*\*\** and *\** respectively indicate a P value of less than 0.001 and 0.05.

To understand which developmental zone the RGF1-treated PLT2 and PLT2^C212S^ proteins regulate at the gene expression level, the DEG data was further compared with our previous root developmental zone-specific transcriptome analysis^16^. We found that RGF1-treated gPLT2^C212S^ induced genes expressed in the meristematic zone more robustly and reduced the genes expressed in the elongation zone more sharply compared to the PLT2 wild-type (Extended Data Fig. 6a, b). These results demonstrated that the PLT2^C212S^ protein, which was more highly accumulated with RGF1 treatment, more potently modulated the developmental zone-specific genes. Furthermore, to examine which developmental event PLT2 and PLT2^C212S^ proteins regulate, we conducted a gene ontology (GO) analysis using common positively and negatively regulated DEGs.

In the upregulated DEGs, the GO highly enriched categories were related to cell proliferation and DNA replication (Fig. 4d). In contrast, in the downregulated DEGs, GOs relevant to cell differentiations and secondary cell wall biogenesis were greatly enriched (Fig. 4d). In addition, we used qRT-PCR to validate that genes related to the mitotic cell cycle were upregulated in the PLT2^C212S^ line and genes involved in cell wall biogenesis were downregulated in the PLT2^C212S^ line after 0.3 nM RGF1 supplement (Fig. 4e, f). These data confirmed that the highly accumulated PLT2^C212S^ protein controlled morphological change to a greater extent than the PLT2 protein by modulating the expression of developmentally specific genes.

ROS modulate various protein features via Cys, not only protein stability. The 212^th^ Cys resides in the *AP2/ERF* DNA binding domain of PLT2. Therefore, we questioned whether the 212^th^ Cys only controlled the protein stability or manipulated the DNA binding domain to control the different gene expressions. To investigate this, we compared the up and downregulated DEGs in the *gPLT2-GFP* and *gPLT2^C212S^-GFP* lines. Over 85% of DEGs up or downregulated upon RGF1 treatment in the *gPLT2-GFP*/*plt2* line overlapped with those in the *gPLT2^C212S^-GFP*/*plt2* line (Fig. 4b); moreover, the common DEG genes among them were more profoundly modulated in *gPLT2^C212S^-GFP* than in *gPLT2-GFP*/*plt2*(Fig. 4c). These results suggested that the PLT2^C212S^ protein dose-dependently modulates expressions of similar target genes but does not change the target genes.

Overall, our results demonstrated that the PLT2 protein stability was modulated through the 212^th^ Cys via ROS signals by RGF1 peptide treatment. Furthermore, the accumulated PLT2 protein dose-dependently regulated its downstream genes to expand the meristematic zone size and decelerate the initiation of cell elongation and differentiation (Fig. 5). These results suggest that the 212^th^ Cys plays a pivotal role in forming the concentration gradients of PLT2 by RGF1 via ROS required for root meristem size control.

**Fig. 5.**
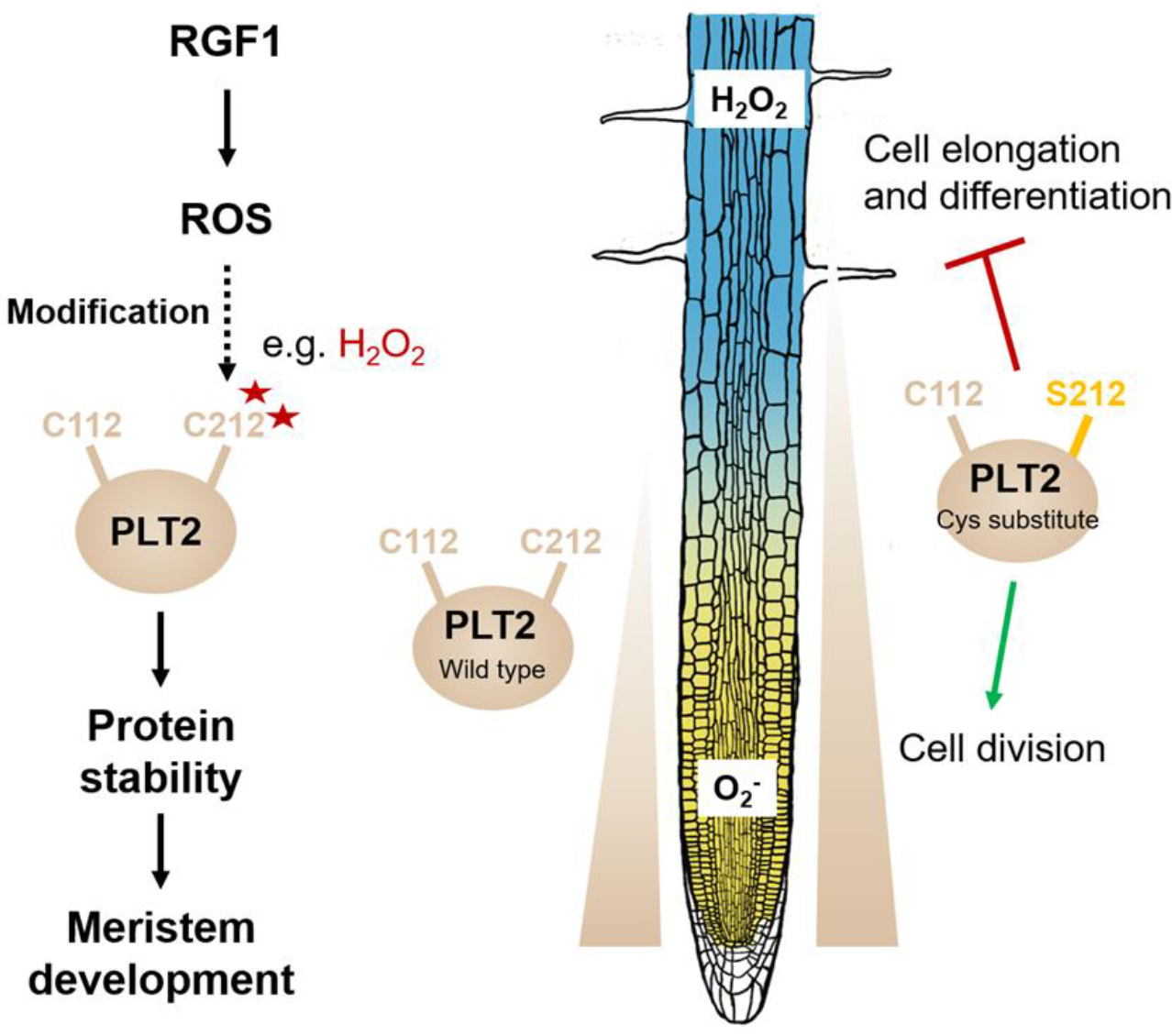
Schema depicting the role of RGF1-ROS in regulating root meristem development via the 212^th^ Cys of PLT2. RGF1 signaling alters ROS distributions along the root developmental zones. The ROS further attacks or triggers a yet-to-be-determined modification on the 212^th^ Cys of PLT2, thereby controlling PLT2 protein stability. In the PLT2 Cys substitute plant, the 212^th^ residue is no longer affected by ROS, so PLT2 shows extended localization. The accumulated PLT2 further induces genes involved in cell proliferation to expand the meristematic zone size and downregulates genes related to cell elongation and differentiation to postpone the initiation of cell maturation. In summary, the 212^th^ Cys is essential in forming the concentration gradients of PLT2 modulated by RGF1-ROS signaling required for root meristem size control.

## Discussion

ROS distributions altered by RGF1 affect the developmental transitions in the root meristem by enhancing PLT2 stability^9^. However, previously, it was unclear how the discrete ROS in each developmental zone affected the PLT2 protein stability. In this study, we provided evidence showing that ROS influence the Cys residues of PLT2, especially the 212^th^, to modulate its stability (Fig. 2, 3). The PLT2^C112S^ protein level was also increased to a greater extent by RGF1 than the wild-type PLT2 but not to the extent of PLT2^C212S^ (Fig. 2c, d, and f). However, the *gPLT2^C112S^-GFP/plt2* line had comparable root meristem phenotypes to the *gPLT2-GFP/plt2* line upon RGF1 treatment (Fig. 2a, b). Previous studies suggested that the PLT protein level has a threshold that controls root development^4,13^. The PLT^C112S^ protein accumulation level may be lower than the threshold level to increase the meristematic zone size. Furthermore, the PLT2^C112S^ protein showed a modest response to H_2_O_2_ (Fig. 3c-g). However, the PLT2^C212S^ protein was almost insensitive to H_2_O_2_ (Fig. 3c-g). These data suggested that the 212^th^ Cys of PLT2 had a more pivotal role in root meristem size regulation under the RGF1 signal via ROS than the 112^th^ Cys.

The 212^th^ Cys is conserved in the DNA binding domain of PLT1 and PLT2 (Extended Data Fig. 1). Even so, only PLT2 showed a significant effect in response to RGF1 via ROS (Fig. 1-3). RGF1 significantly decreases H_2_O_2_ levels around the boundary between the meristematic and elongation zones^9^. The PLT2 concentration gradient is formed until the region where RGF1 affects the ROS distributions (Fig. 3a, b). However, the PLT1 protein distributions are limited around the RAM (Fig. 1e), suggesting that the alterations of H_2_O_2_ levels by RGF1 are not coincident with the existence of the PLT1 protein and do not affect PLT1 distributions. Furthermore, the *PLT1* expression levels were significantly reduced in the *gPLT2-GFP/plt2* line and *gPLT2* ^C212S^ *-GFP/plt2* lines after RGF1 treatment compared with the Mock (Extended Data Fig. 7). In contrast, the *PLT1* expression was comparable in the *plt2* mutant even after RGF1 treatment relative to Mock treatment suggesting that the PLT2 accumulated by RGF1 transcriptionally reduced the *PLT1* expression. These results indicated that RGF1 transcriptionally decreased the PLT1 expression via PLT2 but did not enhance the PLT1 allocation through post-translational regulation via ROS. We thus found that the PLT2 has a distinct function under the RGF1 signaling pathway.

The wild-type PLT2 protein was generated around the RAM and gradually broken down all the way up to the elongation zone, where H_2_O_2_ starts accumulating^9,15^ (Fig. 3a, b). We could detect weak signals at the elongation zone in the *gPLT2-GFP/plt2* line relative to those around RAM. However, the PLT2^C212S^ protein was more slowly degraded and still detected in the elongation zone at a similar level to the proximal side of the meristematic zone (Fig. 3a, b). The PLT2^C212S^ protein demonstrated a robust resistance to a high dose of H_2_O_2_ (Fig. 3e-g) and moderate insensitivity to lower O_2_^-^ (Extended Data Fig. 5a, b). RGF1 treatment strongly reduces H_2_O_2_ levels at the boundary between the meristematic and elongation zones and increases O_2_^-^ concentrations in the meristematic zone^9^. These findings suggest that the 212^th^ Cys is substantially affected by local ROS alteration at specific root sites, presumably by ROS-dependent modification to expand the concentration gradients. Furthermore, we can expect that the PLT2 protein around the transition zone encounters the ROS altered by RGF1 and is modified by them. Therefore, the portions of the modified PLT2 by ROS may be low. Detection of the modified PLT2 proteins is technically tricky. We detected only a few peptides, including the 212^th^ Cys, and could not detect any modification. Further experiments may uncover the specific modification of the 212^th^ Cys.

The distinct localizations of two ROS in individual developmental zones modulate root development as signaling molecules^15,17^. ROS alterations affect various protein activities and stabilities. In this study, we focused only on PLT2. However, ROS are expected to post-translationally regulate many proteins in specific developmental zones. Further cysteine proteome analysis in individual developmental zones may uncover novel target proteins that function via Cys through ROS required for root development in a specific root developmental zone. In summary, our current study provided insight into the post-translational regulation, via the cysteine by ROS, of one of the pivotal root meristem regulators in a specific root developmental zone.

## Methods

### Plant materials and growth conditions

All *Arabidopsis* mutants and transgenic lines used in this study were in the Columbia-0 (Col-0) background. The transfer (T)-DNA insertion line SAIL_846_C09 (*plt1*) was obtained from the Arabidopsis Biological Resource Center at Ohio State University.

SALK_130119.20.25 (*plt2*) and SALK_127417.55.25 (*plt3*) were previously described^4,9^. Seeds were surface-sterilized in 50% bleach coupled with 0.1% Tween 20 (Sigma) for 10 min, then rinsed 5 times with sterile water, followed by stratification at 4°C for 2 days.

The sterilized seeds were plated on half-strength MS medium (Murashige and Skoog salt mixture; Caisson Laboratories) containing 0.05% MES, 1% sucrose, and 1% agar (SIGMA), and that pH was adjusted to 5.7 with KOH. Seedlings were grown vertically in a growth incubator at 22°C with 16 h light/8 h darkness cycles for 7 days. Then, seedlings were transferred to ½ MS agar plates containing either water (mock treatment), synthetic sulfated RGF1 peptide (Mission Biotech, Taiwan), or the indicated chemicals for 24 h for further experimentation.

### Plasmid constructions and transformations into plants

The genomic sequence of the *PLT2* gene, including a native promoter (2410 bp) without a stop codon, was amplified using CloneAmp™ HiFi PCR Premix (Takara Bio, USA) from genomic DNA and then sub-cloned into the *pENTR/D/TOPO* vector (Invitrogen). The following primers were used for g*PLT2* amplification: pENTR gPLT2-F and pENTR gPLT2 Nostop-R (all primer names and sequences are listed in Supplementary Table 2 in this research). The sequence of the *PLT2* gene in the *pENTR/D/TOPO* vector (gPLT2 pENTR) was confirmed by Sanger sequencing.

To generate *gPLT2^C112S^*, *gPLT2^C212S^*, and *gPLT2^C112SC212S^*, site-directed mutagenesis of g*PLT2* was performed using the in-fusion HD cloning Kit (Takara Bio). The further reactions followed the in-fusion HD cloning Kit manual (Takara Bio). Two sets of forward and reverse primers for *gPLT2^C112S^*, *gPLT2^C212S^*, and three set primers were constructed to introduce mutations at Cys112, Cys212, and Cys112Cys212. *gPLT2^C112S^* fragment1: In-fusion gPLT2 Forward new and PLT2 C112S-r.

*gPLT2 ^C112S^*fragment2: PLT2 C112S-f and In-fusion gPLT2 Reverse new. *gPLT2 ^C212S^* fragment1: In-fusion gPLT2 Forward new and PLT2 C212S-r. *gPLT2 ^C212S^* fragment2: PLT2 C212S-f and In-fusion gPLT2 Reverse new. *gPLT2^C112SC212S^* fragment1: In-fusion gPLT2 Forward new and PLT2 C112S-r.

*gPLT2^C112SC212S^* fragment2: PLT2 C112S-f and PLT2 C212S-r. *gPLT2^C112SC212S^* fragment3: PLT2 C212S-f and In-fusion gPLT2 Reverse new.

To generate the linearized pENTR vector from the pENTR vector, inverse PCR was conducted using the following primers: inverse pENTR F and inverse pENTR R.

Two or three DNA fragments were fused to the linearized pENTR vector, confirming the sequences.

The clones of *gPLT2*, *gPLT2^C112S^*, *gPLT2^C212S^*, and *gPLT2^C112SC212S^* were recombined with the *pGWB450*^18^ vector using Gateway™ LR Clonase II (Invitrogen). The constructs of *gPLT2-GFP*, *gPLT2^C112S^-GFP*, *gPLT2^C212S^-GFP*, and *gPLT2^C112S^ ^C212S^-GFP* were transformed into the *plt2* mutant.

The *gPLT2-3xGFP* construct was made based on the *gPLT2-GFP* backbone by replacing the *GFP* with three copies of the *GFP* through infusion cloning. The constructs of *gPLT2-GFP*, gPLT2^C112S^-GFP, *gPLT2^C212S^-GFP*, *gPLT2^C112S^ ^C212S^-GFP*, and *gPLT2-3xGFP* were transformed into *Agrobacterium tumefaciens* GV3101 strain and introduced into *Arabidopsis* (*plt2* background) using the floral dipping method^19^. Transgenic plants were selected on 50 μg/mL kanamycin plates, and the two independent T3 homozygous lines from each genotype were used in this study. The *gPLT1-YFP* and *gPLT2-YFP* lines have been previously generated^4^.

### Microscopic analysis

The seedlings were stained with 200 μM NBT dissolved in 20 mM phosphate buffer (pH 6.1) in the dark for 2 min and rinsed twice with distilled water^9^. Root images were taken using a ×10 objective with a Zeiss AXIO Scope A1.

The GFP or YFP signals were observed using a ×20 objective with a Zeiss LSM 980 laser scanning confocal microscope. Excitation and detection parameters were set as follows: GFP, excitation at 488 nm and detection at 491–544 nm; YFP, excitation at 514 nm and detection at 517–544 nm; PI staining, excitation at 561 nm and detection at 565–747 nm. The microscopic images were stitched and analyzed using the Fiji package of ImageJ^12,20^.

### Measurement of the GFP signals at the specific sites in the developmental zones

We grew the seedlings of the *gPLT2-GFP/plt2* and *gPLT2^C212S^-GFP/plt2* lines for 7 days on ½ MS agar plates. The single-layer images for GFP and PI were taken under the confocal laser scanning microscope (Zeiss LSM 980). The sites close to the apical meristem, near the elongation zone in the meristematic zone, and the elongation zone were identified as the apical meristem, the proximal side of the meristematic zone, and the elongation zone. The GFP intensities of three cells of 28-30 roots in the apical meristem (Fig. 2a, red arrowheads), the proximal side of the meristematic zone (Fig. 2a, yellow arrowheads), and the elongation zone (Fig. 2a, blue arrowheads) were measured with three biological replicates in the *gPLT2-GFP/plt2* and *gPLT2^C212S^-GFP/plt2* lines using the Fiji^12,20^.

### Immunoblotting

Root tip tissues were harvested under a dissection microscope (Leica M205 FCA) using an ophthalmic scalpel (Feather) and homogenized in 1xSDS sample buffer containing 50 mM Tris-HCl (pH 6.8), 10% glycerol, 2% SDS, and 5% β-mercaptoethanol. The protein samples were incubated at 95°C for 10 min. The samples were separated by 10%

SDS-PAGE and transferred to 0.45 μm nitrocellulose membrane. The PLT2-GFP fusion protein abundance was detected by using anti-GFP (cat#632381, 1:3,000 dilution; Clontech) and anti-ACTIN (cat#MA1-744, 1:5,000 dilution; Thermo Fisher) as an internal control. The goat anti-mouse IgG-HRP conjugate (cat#1706516, 1:4,000 dilution; BIO-RAD) and the secondary antibody were used to detect anti-GFP and anti-ACTIN.

The chemiluminescent signals were monitored by a ChemiDoc XRS+ System (BIO-RAD).

### RNA extraction and real-time qPCR analysis

The total RNA was isolated using the RNeasy Micro Kit (Qiagen). Briefly, the root tip tissues were homogenized in extraction buffer and processed following the manufacturer’s protocol. The concentration of total RNA was measured by a Qubit (Invitrogen) instrument. Reverse transcription was performed using SuperScript IV Reverse Transcriptase (Invitrogen). The SYBR Green Supermix (BIO-RAD) was used with a CFX Connect Real-Time PCR Detection System (BIO-RAD) for qPCR. Relative transcript levels were determined by normalizing with *PP2AA3* (At1g13320). Three biological replicates and technical replicates were used for each experiment.

### RNA-seq library preparation and data processing

RNA quality was examined using a 2100 Bioanalyzer (Agilent). The RNA integrity number was above 8.0 in all samples. RNA-seq libraries were generated from 100 ng total RNAs using Universal RNA-Seq with NuQuant library preparation kit (Tecan Trading AG) following the manufacturer’s protocols. The libraries for 5 to 6 biological replicates of mock- and RGF1-treated root samples were sequenced on an Illumina NextSeq 2000 (100 base paired-end reads). The bulk RNA-sequencing reads were aligned to the *Arabidopsis* reference genome TAIR10 using HISAT2^21^, and the mappability for each sample can be found in Supplementary Table 1. The mRNA read counts of genes were computed using featureCounts^22^. Genes expressed (read counts > 0) in at least ten samples are included for analysis. Normalization for sequencing depth and differential gene analysis was performed using DESeq2^23^. Genes with FDR < 5% were considered differentially expressed. Gene ontology analysis was performed using the Bioconductor package topGO^24^. All DEGs for bulk RNA sequencing can be found in Supplementary Data Set 1.

### Statistical analysis and reproducibility

Experiments were independently repeated three times with similar results. No power analysis was done to estimate the sample size. All statistical analyses were performed using R version 4.3.2 (http://www.r-project.org/). For pairwise comparison, data were analyzed with a two-tailed t-test. For multiple comparisons, a Welch’s analysis of variance (ANOVA) with Tukey post hoc test was performed. In the boxplots, the circles represent each individual data point, the horizontal line within boxes represents the median, the upper and lower hinges, respectively, indicate the 75^th^ and 25^th^ percentiles (interquartile range, IQR), and the whiskers show the 1.5 extension of the IQR. Data points out of the whiskers were considered outliers.

## Data availability

RNA-seq data have been deposited at GEO: GSE244175 and are publicly available as of publication.

For reviewer To review GEO accession GSE244175: Go to https://www.ncbi.nlm.nih.gov/geo/query/acc.cgi?acc=GSE244175 Enter token ofknccasttmbjoh into the box.

## Acknowledgments

We thank Miranda Loney and Hironaka Tsukagoshi for comments on the manuscript; Yi-Han Weng for taking confocal images; Renze Heidstra for *gPLT2-YFP*, *gPLT1-YFP*, and *plt3* seeds; the High Throughput Sequencing Core in the Biodiversity Research Center at Academia Sinica for sequencing the Illumina libraries; the Confocal Microscope Core Facility in the Biotechnology Center in Southern Taiwan at Academia Sinica for maintaining the confocal laser scanning microscope; and NGS High Throughput Sequencing Core Facility for maintaining the server at Academia Sinica. This work was funded by the Career Development Award, Academia Sinica, Taiwan (AS-CDA-111-L05), the National Science and Technology Council, Taiwan (112-2311-B-001-021-MY3), (111-2311-B-001-028), and (109-2313-B-001-006-MY3), and the Agricultural Biotechnology Research Center, Academia Sinica, Taiwan to M.Y.

## Author contributions

Y-C.H. and M.Y. conceptualized the study; Y-C.H., S-Y.S., and M.Y. performed all experiments; M-R.Y. and L-K. L. performed the computational and statistical analyses; all the authors wrote the paper.

## Extended Data Figures

**Extended Data Fig. 1.**
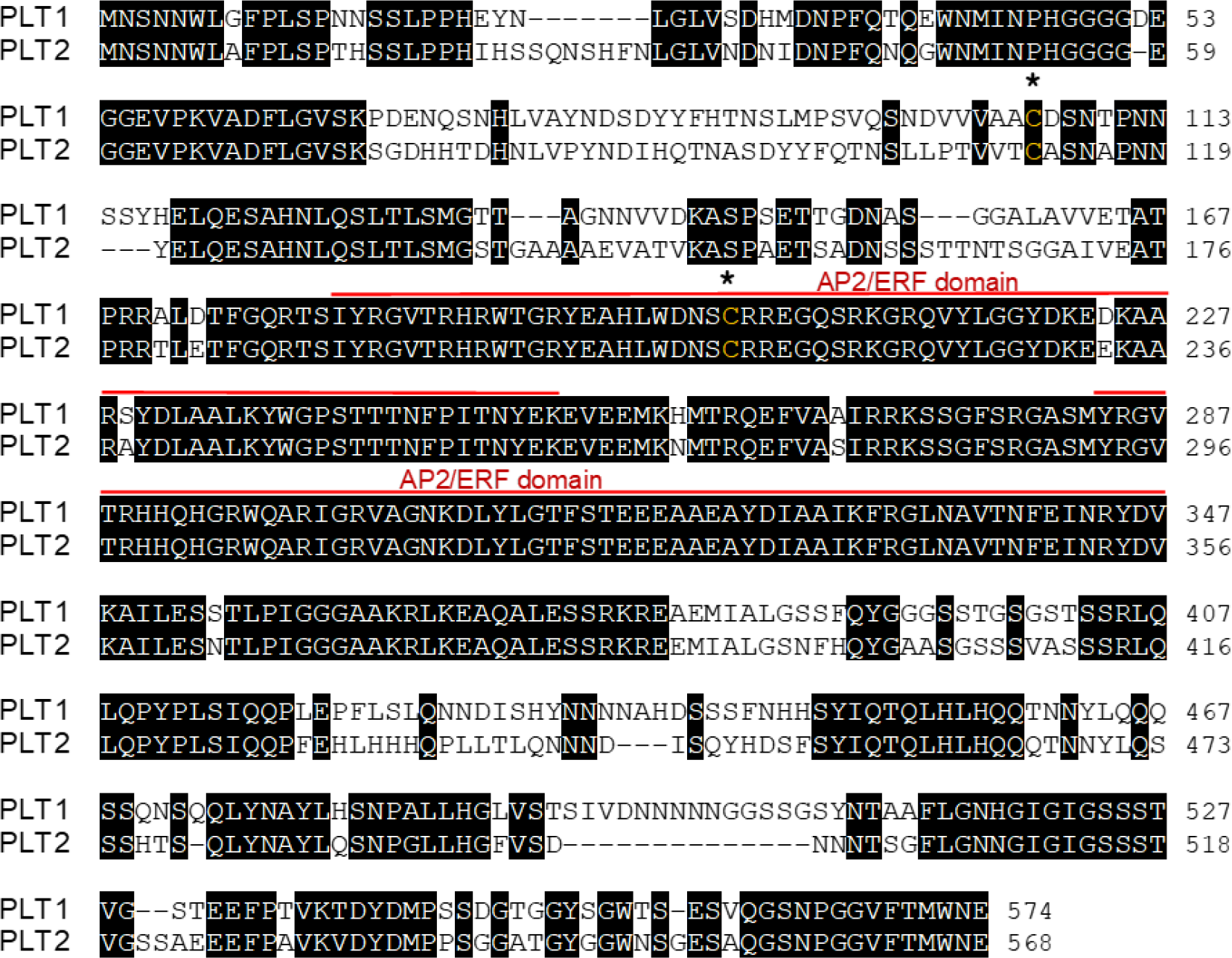
Global sequence alignment of PLT1 and PLT2. Identical amino acid residues are shaded in black. The AP2/ERF domains are indicated with red solid lines above the sequences. Asterisks denote the Cysteine residues conserved in PLT1 and PLT2.

**Extended Data Fig. 2.**
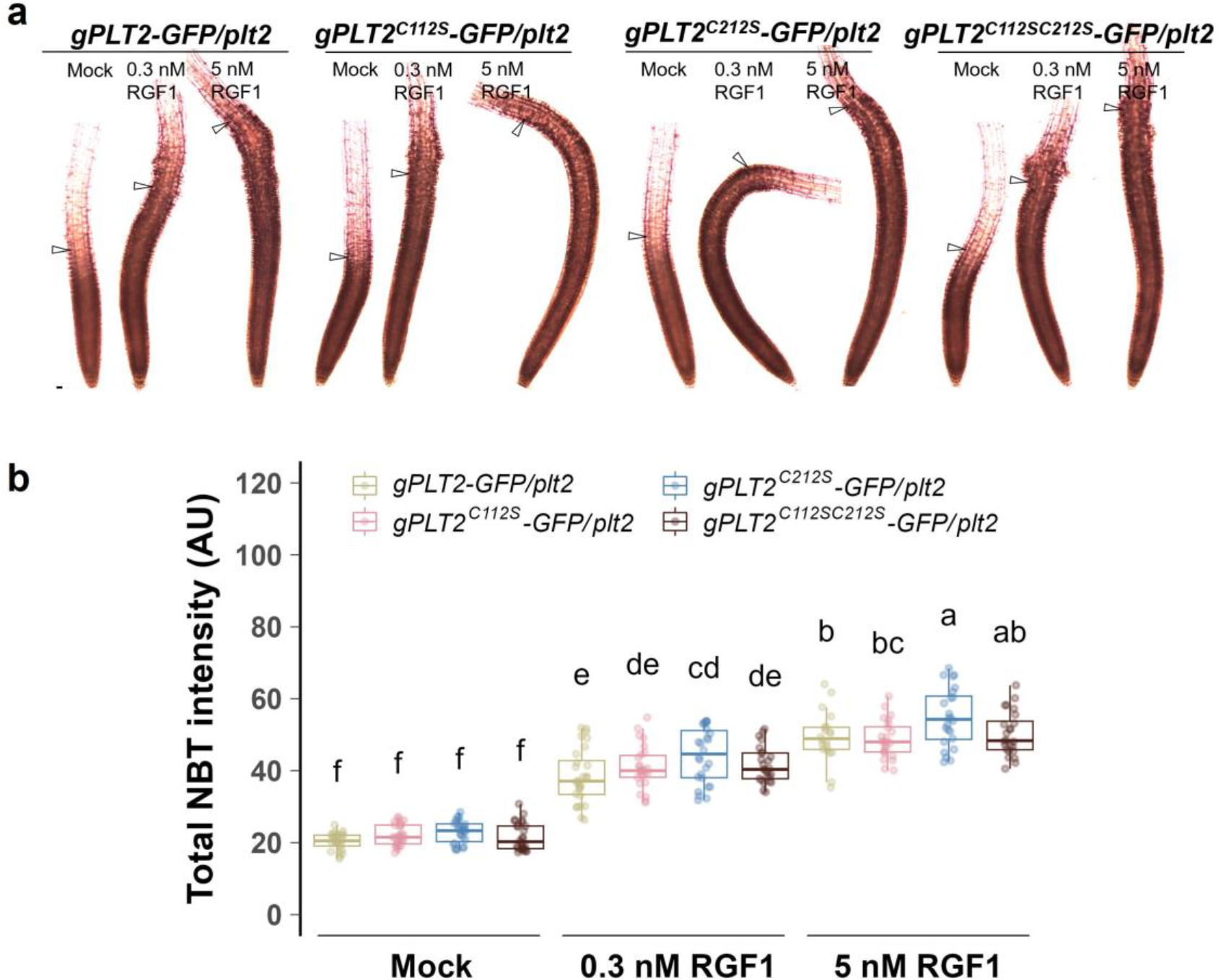
RGF1 treatment similarly modulates O_2_^-^ levels in each PLT2 reporter line. **a**, Light microscope images of *gPLT2-GFP*/*plt2*, *gPLT2^C112S^-GFP*/*plt2*, *gPLT2^C212S^-GFP*/*plt2*, and *gPLT2^C112SC212S^-GFP*/*plt2* roots after mock, 0.3 nM, or 5 nM RGF1 treatment for 24 h followed by NBT staining. The hollow arrowheads indicate the junction between the meristematic and elongation zones. Scale bar, 50 μm. **b**, Quantification of NBT staining intensity in (a). In **b**, the boxplots show the individual data points (circles), median (horizontal line within boxes), interquartile range (IQR) (hinges), and the 1.5 * IQR extension (whiskers). Two-way ANOVA with Tukey’s post hoc test in **b**. Boxes not sharing a lowercase letter indicate a significant difference (*P* < 0.05).

**Extended Data Fig. 3.**
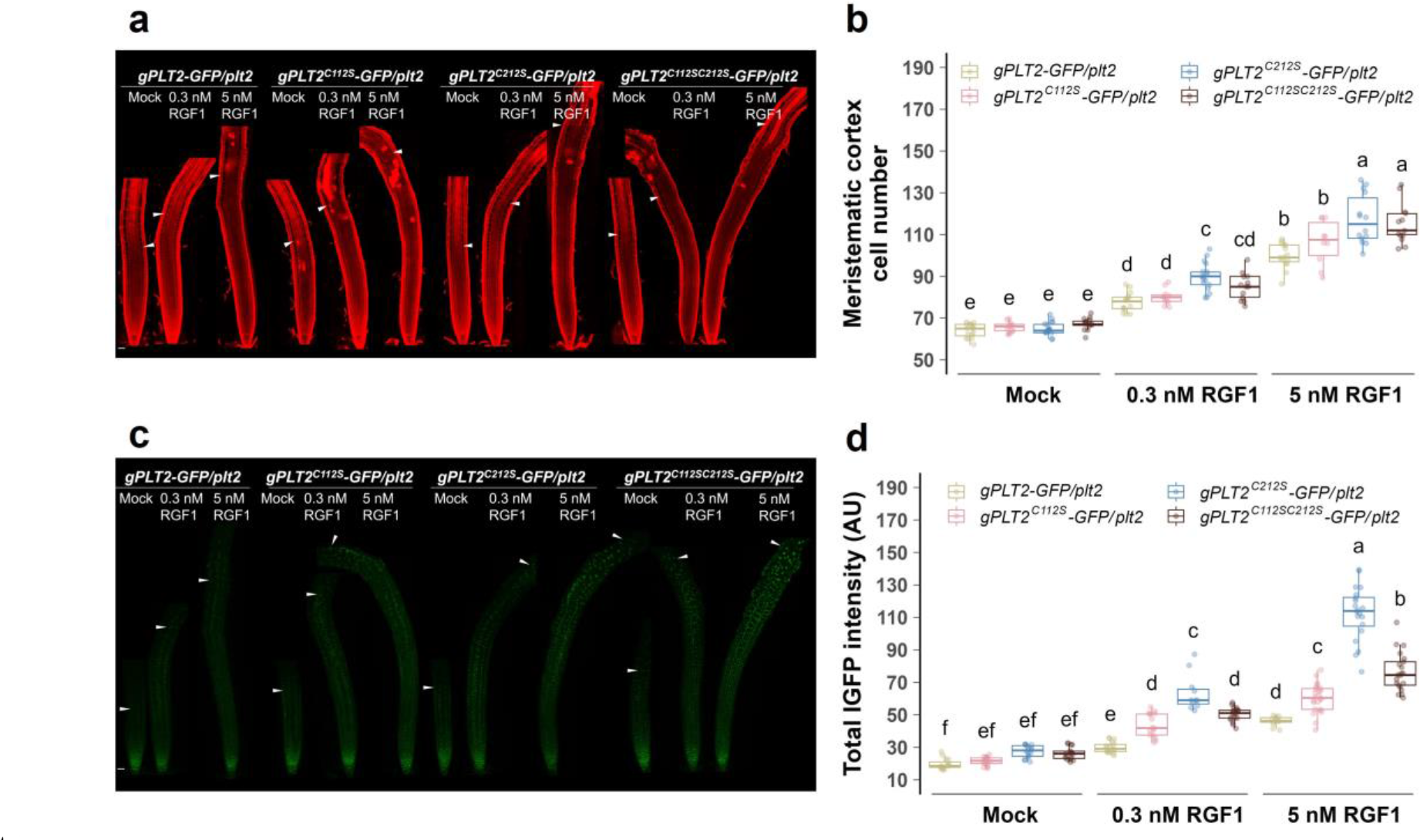
The second independent lines of each PLT2 construct show a similar expression pattern and response to RGF1 treatment. **a**, **b**, **c**, **d**, Confocal images (a and c) and the quantification analysis (b and d) of the second independent line of each *gPLT2-GFP*/*plt2*, *gPLT2^C112S^-GFP*/*plt2*, *gPLT2^C212S^-GFP*/*plt2*, and *gPLT2^C112SC212S^-GFP*/*plt2* roots after mock or 5 nM RGF1 treatment for 24 h. The white arrowheads in a show the junction of the meristematic zones, and in c exhibit the extent of GFP expression. In **b** and **d**, the boxplots show the individual data points (circles), median (horizontal line within boxes), interquartile range (IQR) (hinges), and the 1.5 * IQR extension (whiskers). Two-way ANOVA with Tukey’s post hoc test in **b** and **d**. Boxes not sharing a lower-case letter indicate a significant difference (*P* < 0.05).

**Extended Data Fig. 4.**
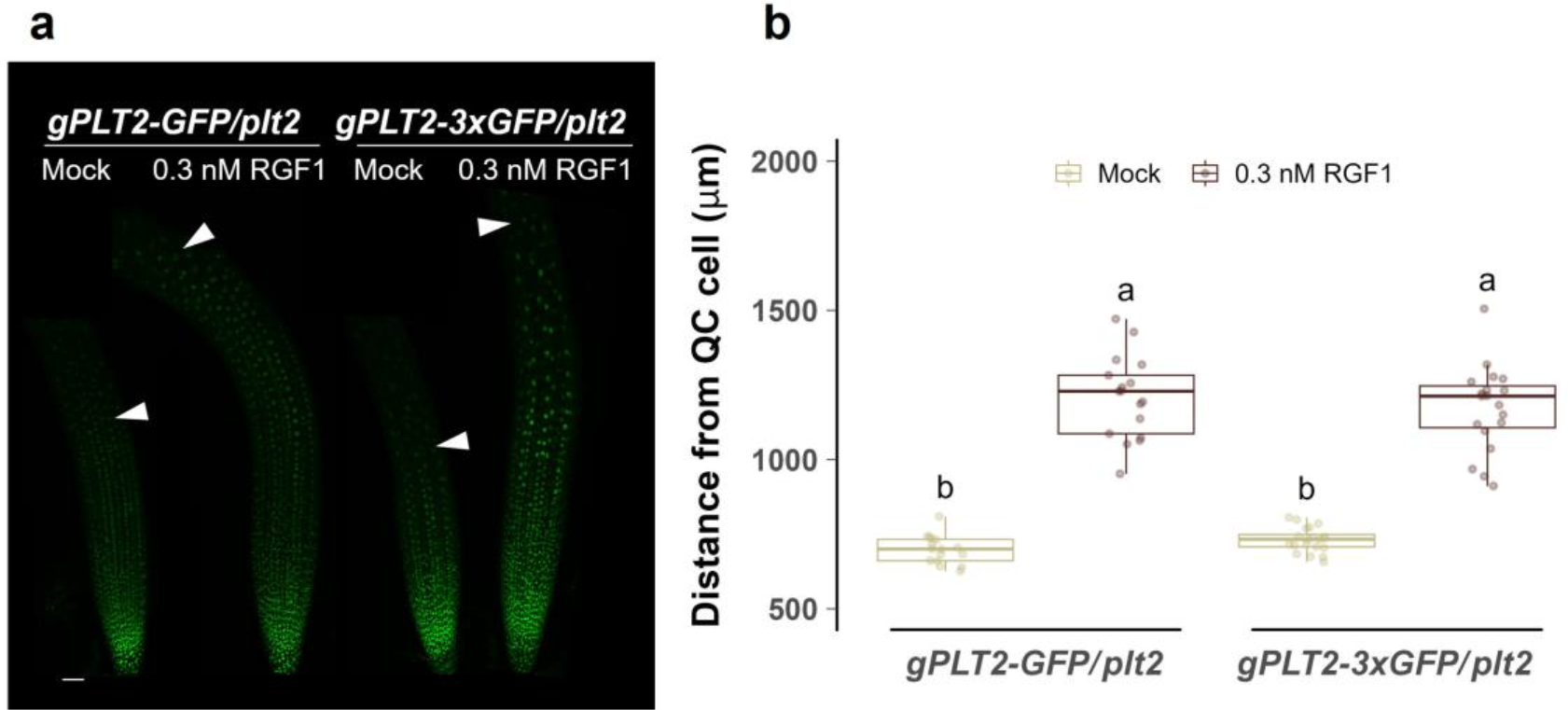
The protein localization of PLT2-1xGFP and PLT2-3xGFP after RGF1 treatment. **a**, Confocal images of the *gPLT2-GFP*/*plt2* and *gPLT2-3xGFP*/*plt2* roots after mock or 0.3 nM RGF1 treatment for 24 h. The white arrowheads indicate the extent of GFP expression. Scale bar, 50 μm. **b**, The GFP quantification of the *gPLT2-GFP*/*plt2* and *gPLT2-3xGFP*/*plt2* roots after mock or 0.3 nM RGF1 treatment for 24 h. In **b**, the boxplots show the individual data points (circles), median (horizontal line within boxes), interquartile range (IQR) (hinges), and the 1.5 * IQR extension (whiskers). Two-way ANOVA with Tukey’s post hoc test in **b**. Boxes not sharing a lowercase letter indicate a significant difference (*P* < 0.05).

**Extended Data Fig. 5.**
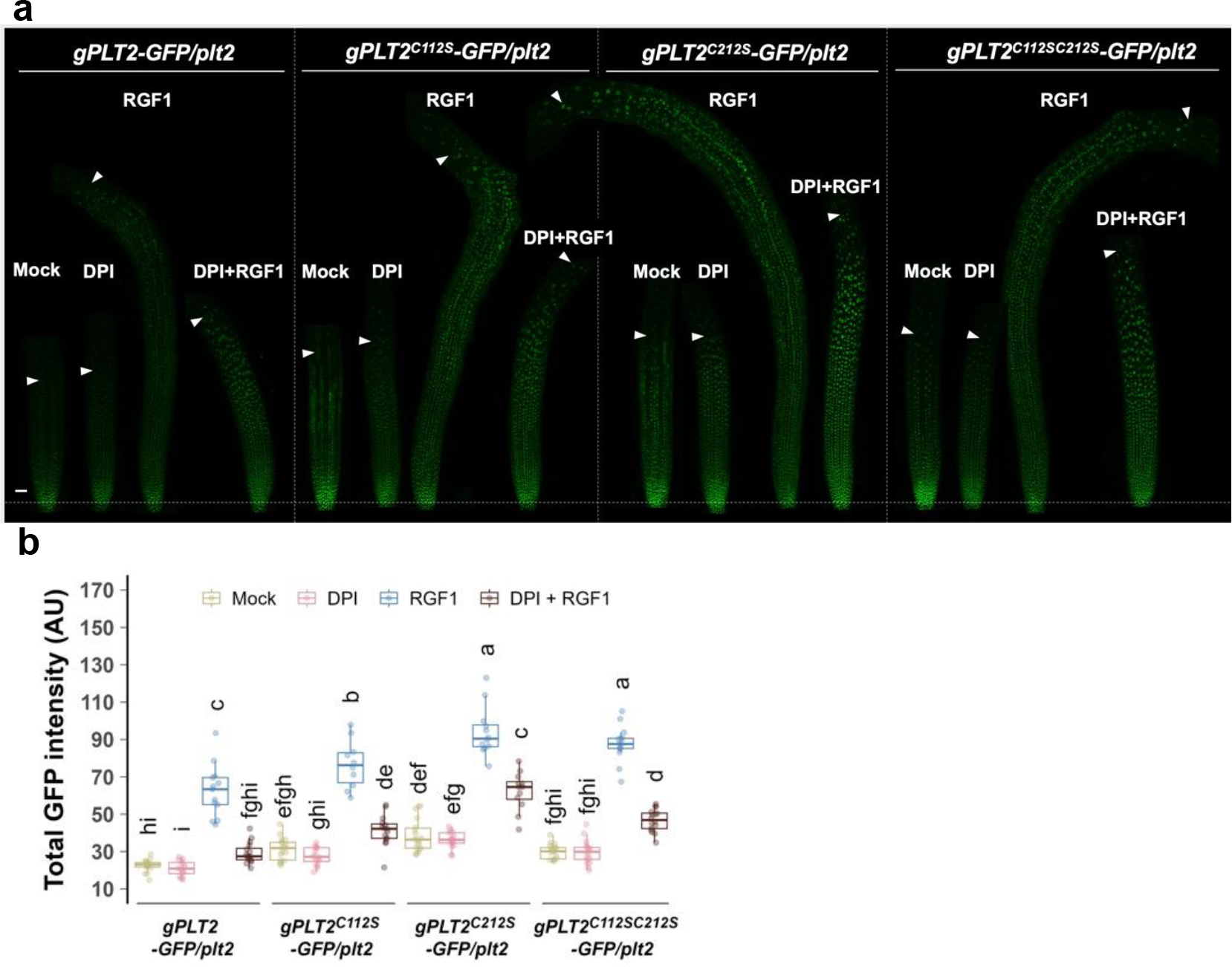
ROS affect the RGF1-dependent PLT2 stability via cysteine residues. **a**, **b**, Confocal images (a) and the quantification analysis (b) of *gPLT2-GFP*/*plt2*, *gPLT2^C112S^-GFP*/*plt2*, *gPLT2^C212S^-GFP*/*plt2*, and *gPLT2^C112SC212S^-GFP*/*plt2* roots were treated with either mock or co-treatment with 10 nM RGF1 and 700 nM DPI (an inhibitor of NADPH oxidase) for 24 h. The white arrowheads in (a) indicate the extent of GFP expression. In **b**, the boxplots show the individual data points (circles), median (horizontal line within boxes), interquartile range (IQR) (hinges), and the 1.5 * IQR extension (whiskers). Two-way ANOVA with Tukey’s post hoc test in **b**. Boxes not sharing a lowercase letter indicate a significant difference (*P* < 0.05). Scale bar, 50 μm (a).

**Extended Data Fig. 6.**
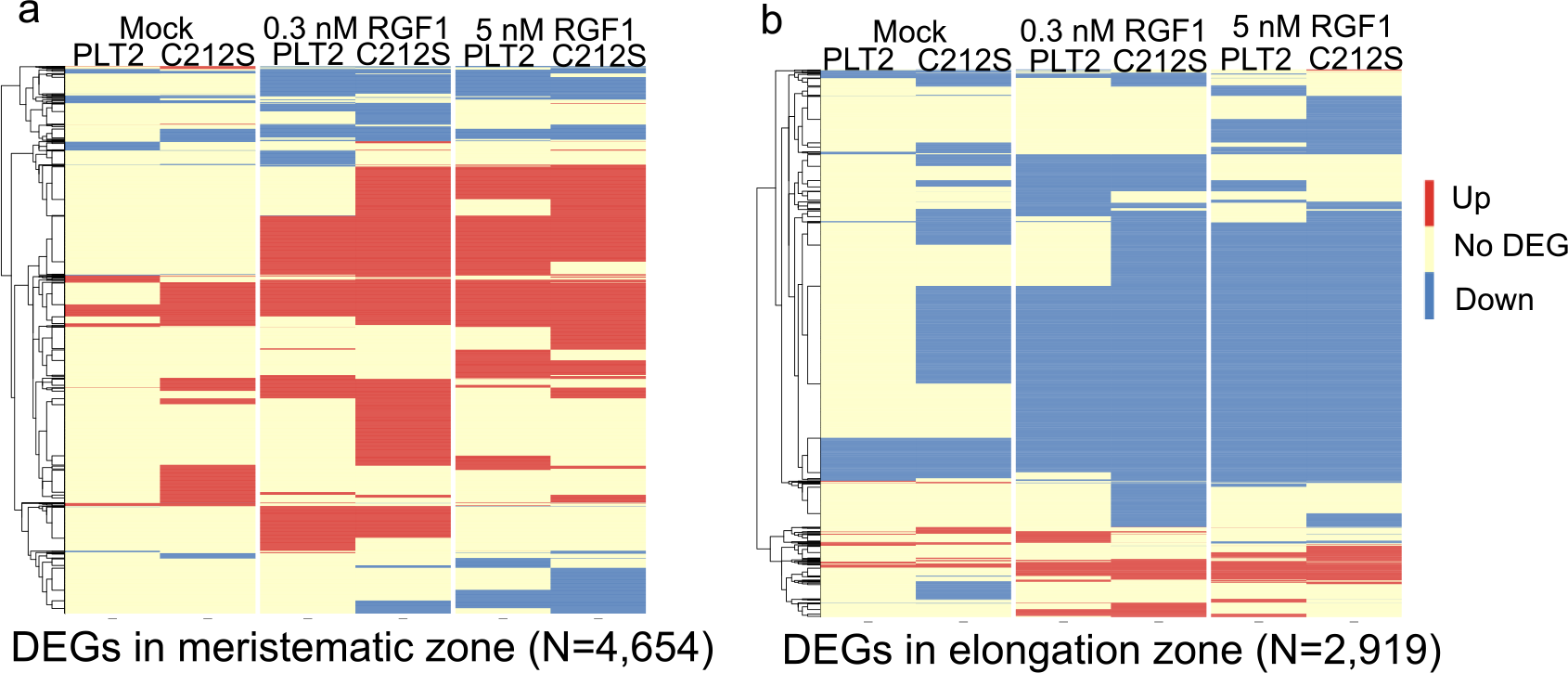
Comparison between all DEGs of PLT2 and PLT2^C212S^ and the specific developmental zone expression genes. **a**, **b**, Expression profiles between upregulated (a) or downregulated (b) DEGs and the meristematic (a) or elongation zone-specific genes (b) according to previous high-resolution root expression map^16^.

**Extended Data Fig. 7.**
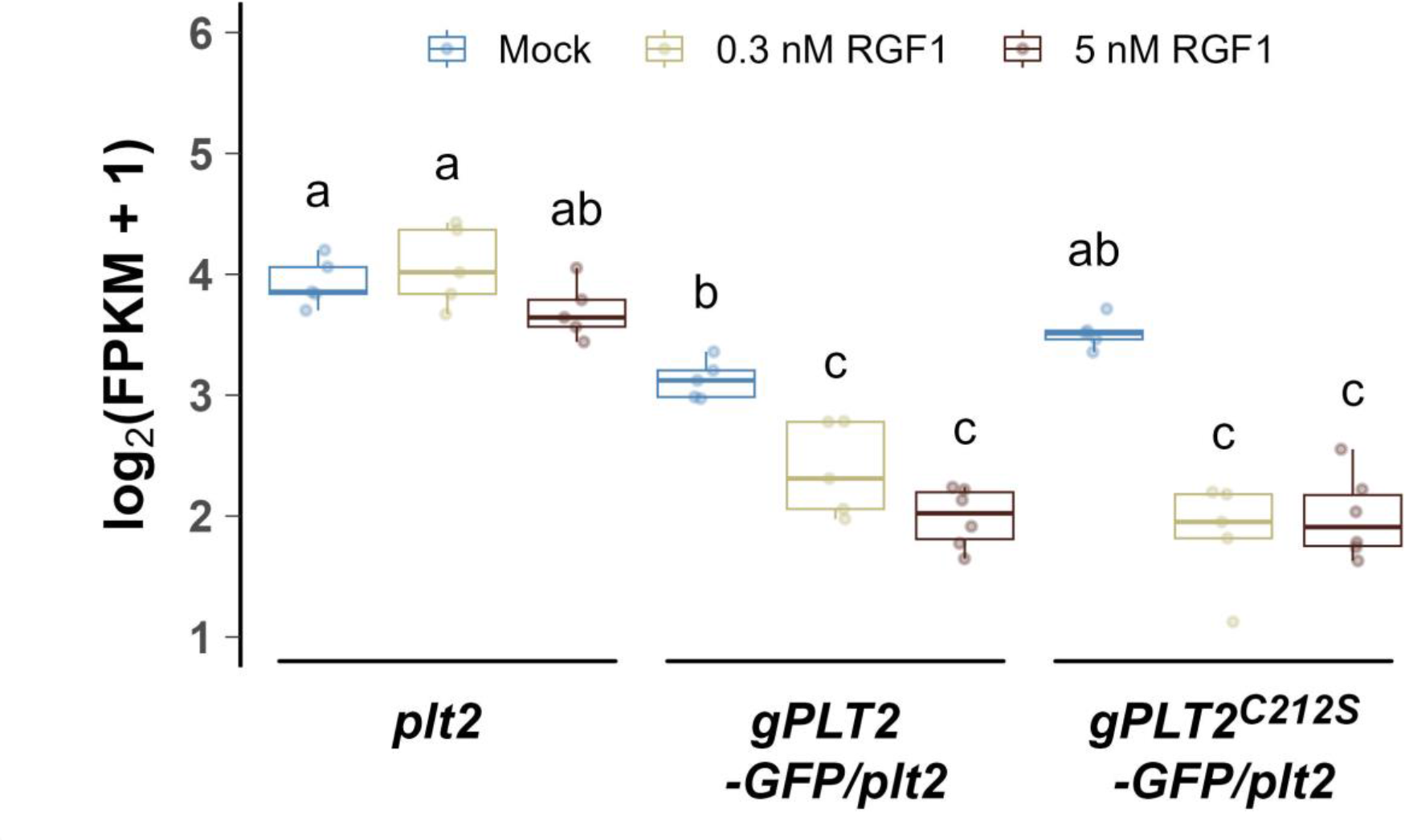
The *PLT1* gene expression in *plt2*, *gPLT2-GFP/plt2* and *gPLT2^C212S^-GFP/plt2* upon RGF1 treatment. The boxplots show the individual data points (circles), median (horizontal line within boxes), interquartile range (IQR) (hinges), and the 1.5 * IQR extension (whiskers). Two-way ANOVA with Tukey’s post hoc test. Boxes not sharing a lowercase letter indicate a significant difference (*P* < 0.05).

